# Islands promote diversification within the silvereye clade: a phylogenomic analysis of a great speciator

**DOI:** 10.1101/2024.03.11.584438

**Authors:** Andrea Estandía, Nilo Merino Recalde, Ashley T. Sendell-Price, Dominique Potvin, Bruce Robertson, Sonya Clegg

## Abstract

Geographic isolation plays a pivotal role in speciation by restricting gene flow between populations through distance or physical barriers. However, the speciation process is complex, influenced by the interplay between dispersal ability and geographic isolation, especially in “great speciators” – bird species present on multiple islands that, at the same time, have many subspecies. Comparing population differentiation in both continental and insular settings can help us to understand the importance of geographical context in the emergence of great speciators. The highly diverse white-eye family Zosteropidae includes several great speciators, including the silvereye (*Zosterops lateralis*) which consists of 16 subspecies, 11 occurring on islands. The distribution of the silvereye on the Australian continent and numerous southwest Pacific islands allows us to explore the influence of different forms of geographic isolation on population divergence. To do this, we conducted a comprehensive phylogenomic analysis of the silvereye and compared patterns of population divergence in insular versus continental silvereye populations. We estimate that the silvereye lineage emerged approximately 1.5 million years ago, followed by the split of the two main silvereye clades: Southern Melanesia and the broader South Pacific (encompassing Australia, New Zealand, and outlying islands). Continental populations show low genetic population structure, which suggests that they can overcome multiple forms of geographic barriers across long distances. In contrast, most island populations are highly structured even over relatively short distances. Divergence statistics further support the idea that water barriers lead to a higher population differentiation when compared to continental distances. Our results indicate that islands promote divergence and provide an empirical example of the geographical conditions that result in the emergence of great speciators.

## Introduction

Geography plays a key role in the speciation process by limiting allele exchange in distant and isolated populations, resulting in divergence (Anderson & Weir, 2022; Bolnick & Fitzpatrick, 2007; Coyne & Allen Orr, 1998). Spatially separated populations may experience isolation-by-distance whereby individuals that are close to each other are likely to be more related than those farther apart (Wright, 1943). Additionally, geographical barriers, such as mountain ranges, deserts or water barriers, can further promote divergence (White, 2016). Oceanic islands and archipelagos are hotspots of biodiversity and home to evolutionary radiations (Miles et al., 2023), thanks in part to their isolation, which acts as a geographical barrier that limits gene flow (Emerson, 2002). Highly dispersive species can colonise distant islands and, by virtue of this vagility, continue to connect populations through gene flow resulting in little population structure (Andersen et al., 2015; Bohonak, 1999; Suárez et al., 2022). However, some taxa, the so-called ‘great speciators’, represent a seemingly paradoxical case. These bird species are present on multiple islands, which indicates high dispersal ability, while having many subspecies, which requires reduced gene flow (Diamond et al., 1976).

Great speciators can be thought of as species with multiple differentiated populations sampled at a particular point along the speciation continuum. Under the ‘taxon cycle’ framework—the idea that speciation on islands proceeds through phases of range expansion and contraction (Wilson, 1961) —a great speciator has undergone the first two phases of a cycle: an initial expansion followed by a reduction in dispersive behaviour that promotes population differentiation and sub-speciation. This pattern can be seen in multiple south-west Pacific birds, including kingfishers (Andersen et al., 2018; DeRaad et al., 2023; O’Connell et al., 2019), corvids (Pepke et al., 2019) and passerines (Estandía, Sendell-Price, Oatley, et al., 2023; Klicka et al., 2023; Pedersen et al., 2018).

Water barriers can be more effective at driving population differentiation compared to continental distances (Broyles et al., 2023; Estandía & Merino Recalde, 2024). Comparing divergence patterns between continental and insular settings can help us to understand the importance of the geographical context in the emergence of great speciators. A fundamental first step in doing this is to have a good understanding of their evolutionary history. Traditionally, phenotypic traits were used to reconstruct phylogenetic relationships, but island taxa often show repeated morphological evolution (Benítez-López et al., 2021), which might lead to an incorrect topology. Genomics now allows us to reconstruct evolutionary relationships with increased confidence (Lee & Palci, 2015), although it is still challenging for those clades that radiate rapidly, such as many great speciators (Moyle et al., 2009). The condensed sequence of cladogenetic events and increased levels of incomplete lineage sorting (ILS) driven by rapid radiation (DeRaad et al., 2023) and recent divergence (Irestedt et al., 2013) complicate phylogenetic inference (Meleshko et al., 2021). Additionally, gene flow can lead to reticulate evolution (Xu, 2000). Conflictive nodes can be identified by using a combination of multiple phylogenetic approaches (Smith et al., 2020) and dense sampling across the genome can provide reliable phylogenetic reconstructions if sources of error are appropriately handled (Kapli et al., 2020).

The white-eye family, Zosteropidae, which originated in southeast Asia (Gwee et al., 2020), has undergone a rapid expansion into the Palaeotropics and Oceania within the last 2 million years (Moyle et al., 2009), giving rise to over 120 species (Clements Checklist v2021). In their studies of northern Melanesian avifauna, Mayr & Diamond (2001) identified *Zosterops griseotinctus* as a great speciator based on the number and distribution of its morphological subspecies. *Zosterops* species outside of this region can also be categorised as great speciators, including the southwestern Pacific silvereye (*Zosterops lateralis*). This species has at least 16 morphological subspecies (11 on islands and the rest on the Australian continent) (Mees, 1969). The island populations have been hypothesised to represent multiple independent colonisations from the Australian mainland (Mees, 1969), with the most recent colonisation from Tasmania to New Zealand and outlying islands being historically documented (Clegg et al., 2002; Sendell-Price et al., 2021).

On the Australian mainland, five subspecies of silvereye are currently recognised based on morphology and plumage patterns (Mees, 1969; Schodde & Mason, 1999). Banding records show they can disperse long distances (Fig. S1) and two subspecies display partial migration: members of the Tasmanian subspecies (*Z. l. lateralis*) and a southern mainland subspecies present in Victoria and New South Wales (*Z. l. westernensis*) journey up to 1,500 km north in the winter. There is evidence of substantial gene flow among eastern Australian subspecies (Estandía, Sendell-Price, Oatley, et al., 2023), but relationships with other mainland subspecies are unclear. Those from Western Australia (*Z. l. chloronotus*) and South Australia (*Z. l. pinarochrous*) are separated from each other and eastern Australian subspecies by major biogeographical barriers. These barriers include the Nullarbor Plain, a desert that stretches approximately 1000 kilometres between Western and South Australia, and the Eyrean barrier, a historical biogeographical barrier inferred to have been present in South Australia. Although the Nullarbor Plain covers a vast area, it has had a lesser impact on current avian diversity compared to the Eyrean barrier (Dolman & Joseph, 2015).

Islands might promote diversification in silvereye populations relative to their continental counterparts. To explore the role of geography in driving divergence in this great speciator, we sequenced the whole genomes of 114 individuals from 8 different white-eye species, including 12 silvereye subspecies from islands and the Australian continent. We explored the phylogenomic relationships among the silvereye complex and related white-eye species to i) reconstruct the evolutionary history of the silvereye; which then provides the basis to ii) compare connectivity patterns within continental versus island subspecies; and iii) determine the role of geography in promoting population differentiation and incipient speciation.

## Methods

### Field sampling

We sampled seven South Pacific white-eye species (the silvereye *Z. lateralis*, representing 12 subspecies, the Vanuatu white-eye *Z. flavifrons*, the Louisiade white-eye *Z. griseotinctus*, the large Lifou white-eye *Z. inornatus*, the small Lifou white-eye *Z. minutus*, the slender-billed white-eye *Z. tenuirostris*, and the green-backed white-eye *Z. xanthochroa*) and the Réunion grey white-eye *Z. borbonicus*, which served as an outgroup (Table S1). Birds were caught in the wild using mist nets or hand-operated traps. A blood sample of 20–40 µl was collected from the brachial wing vein and stored in 1 ml of Queen’s lysis buffer (Seutin et al., 1991). We took a suite of morphological measurements: wing length (maximum flattened chord of the longest primary feather measured with a metal ruler), tail length (the length of the central tail feathers from the base to the tip measured with dividers); metatarsal length, head length from the rear of the skull to the bill tip, and bill length, width, and depth at the posterior nostril opening, all measured with dial callipers.

### DNA extraction, library preparation, and sequencing

We extracted DNA from blood samples using a standard phenol-chloroform protocol (Seutin et al., 1991). We added 100 µl of blood sample to a microcentrifuge tube containing 250 µl of DIGSOL extraction buffer (0.02 M EDTA, 0.05 M Tris-HCl (pH 8.0), 0.4 M NaCl, 0.5% sodium dodecyl sulphate (SDS)) and 10 µl of Proteinase K (20mg/mL). Samples were incubated at 55°C overnight. Following incubation, we added 250 µl of phenol:chloroform:isoamyl alcohol (25:24:1) and gently mixed the samples for 10 minutes, then centrifuged at 10,000 rpm for 10 minutes. We transferred the aqueous layer to a new microcentrifuge tube and repeated the previous step by adding the phenol mixture, mixing and centrifuging it. We recovered the aqueous phase, transferred it to a new tube, added 250 µl of chloroform:isolamyl alcohol (24:1), and mixed and centrifuged the samples as in the previous steps. We precipitated the DNA by adding 2 volumes of ice-cold 100% ethanol, 1 volume of 2.5M ammonium acetate, and 2 µl of glycogen, leaving it overnight at -20°C, then centrifuging the tubes for 10 min at 15,000 rpm at 4°C before removing the supernatant. We rinsed the precipitate with 500 µl of ice-cold 70% ethanol and centrifuged the tubes as in the previous step. We removed the supernatant and left the precipitated DNA to dry at room temperature. Once dried, the DNA was resuspended in 50 µl of TE (Tris-EDTA) buffer (0.01 M Tris-HCL (pH 8.0), 0.0001 M EDTA). We quantified the DNA concentration with a Qubit 2.0. DNA extracts were sent to Novogene UK for library preparation and whole genome sequencing at 5X on the Illumina Novaseq 6000 platform (Illumina, San Diego) using paired-end 150bp sequencing reads.

### Quality control and genotype calling

We trimmed reads to remove adapter content and base calls of low quality using fastp (Chen et al., 2018) with default settings and 10 bp trimmed from the start of each read.

We aligned cleaned reads of each individual to a pseudochromosome assembly for this species (Estandía, Sendell-Price, Robertson, et al., 2023) using the Burrows-Wheeler Aligner (BWA)-mem algorithm (Li & Durbin, 2010). We calculated genotype probabilities in ANGSD (Korneliussen et al., 2014) using the samtools model and keeping SNPs with a minimum allele frequency of 0.05 and a mapping quality score 20. We filtered the bam files by excluding SNPs that showed significant deviation from Hardy-Weinberg equilibrium and excluded any loci putatively under selection with PCAngsd (Meisner & Albrechtsen, 2018) to have an unbiased neutral dataset that allows accurate population structure analyses (Kirk & Freeland, 2011). We further filtered our dataset by only retaining those sites that were placed on autosomes for which genotype probabilities were high (100% confidence), allowing us to call genotypes and obtain a VCF file with 21,726 SNPs suitable for analyses of population structure and phylogenetic reconstruction.

### Phylogenetic reconstruction

We conducted three independent maximum likelihood analyses with IQ-TREE (Nguyen et al., 2015) with the most appropriate model for our data, selected using ModelFinder (Kalyaanamoorthy et al., 2017). As SNPs-based datasets only contain variable sites and can lead to branch length overestimation (Leaché et al., 2015), we applied ascertainment bias correction. We obtained support values by running 100 bootstraps and applying a 25% burn-in.

Concatenated approaches typically do poorly in the presence of ILS (Warnow, 2015). An alternative approach, the multispecies coalescent model, offers a solution by inferring phylogenies while accounting for ancestral polymorphisms and gene tree-species tree conflict (Edwards et al., 2016). We estimated species trees using two coalescent-based methods: SNAPPER, implemented within BEAST2 (Bouckaert et al., 2014); and SVDquartets, implemented within PAUP* (Swofford & Sullivan, 2003) which is a method robust to the presence of gene flow (Long & Kubatko, 2018).

SVDquartets takes multi-locus single-site data, infers the quartet trees for all subsets of four species, and then combines these trees into a species tree (Chou et al., 2015). We used SVDquartets to build 100,000 random quartet trees with the main genomic dataset (21,726 SNPs). We then explored uncertainty in relationships by producing 100 bootstrap replicates. We used Figtree (Rambaut, 2009) to visualise a majority-rule consensus tree, whereby only relationships that appear in at least 50% of rival trees are resolved. Since SNAPPER is more computationally intensive, we randomly selected 1500 SNPs, where all loci were called at high confidence. We set four priors: 1. *Z. borbonicus* was set as an outgroup based on previous white-eye phylogenies (Gwee et al., 2020); 2. the silvereye was set as an independent clade; 3. we time-calibrated the root of the tree by applying a normal distribution with an offset at 2.25 Mya, mean of 0.8 and SD of 1, ensuring that 95% of the probability falls within the timeframe when *Zosterops* emerged (Leroy et al., 2021); and 4. we set a prior for the emergence of the Capricorn silvereye (*Z. l. chlorocephalus*) at 4,000 years old with a SD of 1,000 as the islands that the subspecies currently inhabits were formed approximately 4,400 years ago and have been vegetated for no more than 4000 years (Clegg et al., 2008; Hopley et al., 2007). We ran three independent analyses with a Markov Chain Monte Carlo (MCMC) chain of 100,000 steps with a burn-in of 10,000 steps. We visualised the full set of trees with Densitree (Bouckaert, 2010), highlighting inconsistencies when there were conflicting tree topologies.

To explore the occurrence of reticulate evolution we used Splitstree4 (Huson & Bryant, 2006) and built a Neighbor Net, which visualises genetic distance between individual samples as a phylogenetic network (Bryant & Moulton, 2004). This approach helps represent complex evolutionary histories that do not fit neatly into a bifurcating tree structure.

### Population structure

We performed a genomic PCA with PCAngsd and decomposed the resulting covariance matrix into eigenvectors to explore population structure visually. We also conducted admixture analyses with NGSadmix (Skotte et al., 2013) by setting a maximum of 22 clusters, the total number of species plus silvereye populations sampled. To explore substructure within two highly divergent groups, we created two separate datasets (Dataset 1: Southern Melanesia (SM) and Dataset 2: Australia, New Zealand and Outlying Islands (ANZO)) and repeated the PCA and admixture analyses described above. We used WGSassign (DeSaix et al., 2023) to assign individuals to populations. When populations are genetically distinct, individuals can be assigned to their population of origin with high confidence, whereas highly admixed populations result in low confidence assignments and many individuals are ‘misassigned’ to non-origin populations. We defined populations at the level of island or continental region (State or Territory of Australia, except for the Australian Capital Territory, which was treated as part of New South Wales). We used a leave-one-out approach, removing one sample at a time from the sampled population, recalculating that population’s allele frequency, and then estimating how likely it was for that sample to belong to each of the tested populations. Following DeSaix et al., (2024) we calculated posterior probabilities of assignment to a population by dividing the maximum likelihood of assignment over the sum of all likelihoods.

### Gene flow

In addition to ILS, admixture across highly divergent lineages could lead to phylogenetic conflict. We examined if admixture among mainland species explains unresolved nodes. We restricted this analysis to mainland subspecies because most widely used models assume constant population sizes and no selection (Smith & Hahn, 2023), an assumption that is violated in island silvereye populations that have experienced size fluctuations associated with colonisation and establishment (Estoup & Clegg, 2003). We explored patterns of population splits and mixtures in the history of mainland subspecies with Treemix (Pickrell & Pritchard, 2012), which fits a model based on population allele frequencies and a Gaussian approximation to genetic drift. We generated maximum likelihood trees taking into account linkage disequilibrium by grouping 1000 SNPs into blocks. We tested a range of migration events (m from 0 to 15) and estimated the optimal migration edges with OptM (Fitak, 2021), which resulted in a maximum of four migration events.

### Morphological distinctiveness

We trained a Random Forest Classifier on a dataset of morphological data from 1292 individuals from 21 silvereye populations (Table S2) for which we had measured the following traits: wing, tail, metatarsal and head length, posterior bill length, and anterior bill width and depth. The model training process was divided into three steps:

- Data Resampling: To address the marked class imbalance, we resampled the data with replacement so that each population had the same number of samples. This was done by grouping the data by population and sampling 15 instances from each group. This process was repeated 100 times to get a better estimate of the variance in model performance on random subsets.
- Model Training: We split the resampled data into training and testing sets, trained a Random Forest Classifier on the training data, and made predictions on the testing data. At each iteration, we added the confusion matrix of the predictions to an average confusion matrix and stored the feature importances to quantify uncertainty derived from stochasticity in both model performance and data selection.
- Model Evaluation: Model Evaluation: After training we evaluated the model’s performance on the testing data. This included calculating accuracy, generating a confusion matrix, and performing a permutation test. We generated several plots to visualise the model’s performance, including a permutation test score plot (Fig. S2), a confusion matrix heatmap, a hierarchical clustering of the confusion matrix, and a feature importance plot (Fig. S3).

The goal of this process was to train a model to classify populations based on their morphology, on the assumption that the degree of misclassification between pairs of populations is indicative of their overall morphological (dis)similarity based on a non-linear combination of traits. This, in combination with the molecular data, can inform us of the extent to which genomic divergence is paralleled by morphological changes, and highlight cases of either parallel morphological changes or conservatism.

### Testing the geographical drivers of population differentiation

To explore whether water gaps promote differentiation more than continental distances, we built a model in *brms* (Bürkner, 2017) that incorporated: i) pairwise F_ST_ as the outcome variable, which represents population differentiation and was calculated with ANGSD; ii) the logarithm of distance between each of the populations as a predictor, iii) and whether the comparison was between islands or between continental populations, also as a predictor. We also tested for an interaction between the two predictors and added a multi-membership grouping term. We used MCMC with four chains of 4000 iterations each, including a warm-up of 400 iterations. We evaluated convergence via visual inspection of the MCMC trace plots, checking that the ESS>200 and the R values for each parameter (R=1 at convergence).

## Results

### Phylogenomic analyses

We used several approaches to explore the evolutionary relationships of this group of birds: maximum-likelihood with IQ-TREE (Fig. S4), coalescence model-based SVDquartets with PAUP* (Fig. 1A), and Bayesian inference with SNAPPER (Fig. 1B). All analyses agreed on a split between South Pacific *Zosterops* species and *Z. borbonicus*, which we estimated to have occurred around 2.17 Ma (95% credibility intervals (CI)=1.45-2.9) (Fig. 1; Fig. S5). The Louisiade white-eye *Z. griseotinctus* was the earliest group to branch off, followed by *Z. flavifrons*, and then *Z. tenuirostris*. Among these, the split between *Z. tenuirostris* and the three other non-silvereye species in New Caledonia ( *Z. inornatus, Z. minutus*, and *Z. xanthochroa*) showed different patterns of relatedness depending on the analytical method used: SNAPPER supported *Z. tenuirostris* as sister to the three species of New Caledonian white-eyes (Fig. 1B), IQ-TREE proposed placing it as the sister of the Large Lifou white-eye *Z. inornatus* (Fig. S4), and SVDquartets indicated a polytomy (Fig. 1A). All analyses showed that *Z. minutus* and *Z. xanthochroa* were more closely related than *Z. inornatus* and *Z. minutus* that occur as single-island endemics on Lifou. The silvereye split from other *Zosterops* species approximately 1.5 Ma (95% CI=0.98-2.01), rapidly forming two separate clades representing Southern Melanesia, which split into Vanuatu and New Caledonia, and ANZO (those from Australia, New Zealand, and the Outlying Islands), with Lord Howe Island as the basal lineage of this group. Despite showing high support for the overall divergence pattern at these nodes, the credibility intervals for their ages are wide (Fig. S5). While these deeper relationships were supported by high bootstrap values and posterior probabilities, shallower ones often showed less support. For example, Southern Melanesia showed complex relationships: SVDquartets indicated a split between New Caledonia and Vanuatu (Fig. 1A). However, SNAPPER suggested Tanna as the basal lineage, followed by Mare (Fig. 1B), while IQ-TREE suggested an early split of a clade containing Mare and Grand Terre (Fig. S4).

**Fig. 1.**
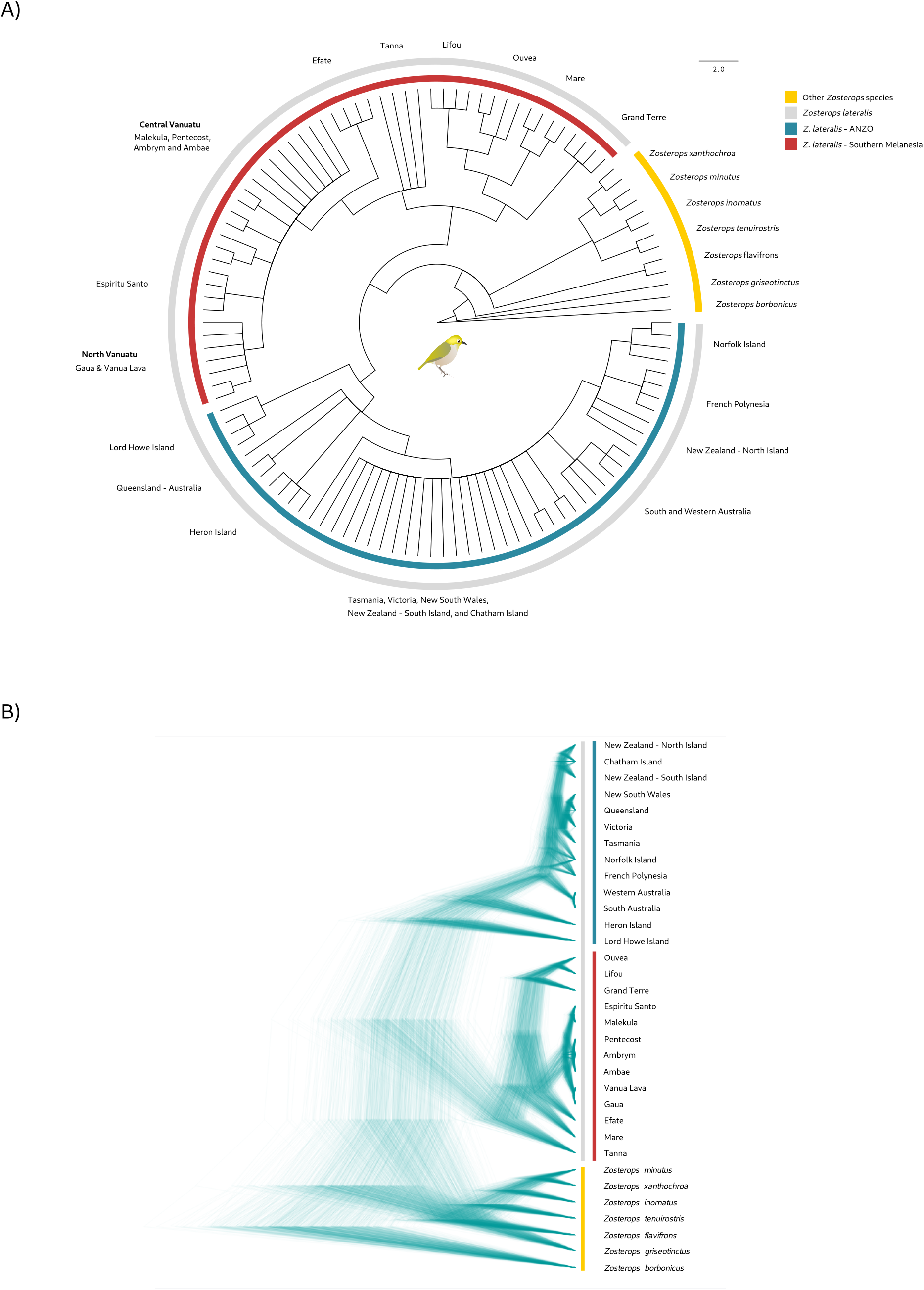
Phylogenomic reconstruction of the silvereye clade and other South Pacific white-eyes. A) Phylogenetic relationships across South Pacific white-eyes, including a dense sampling of silvereye subspecies using a coalescent-based approach with SVDquartets. B) Variability in the inferred phylogenetic relationships with SNAPPER, highlighting areas where the tree space is more densely populated. Phylogenetic conflict occurs at shallow nodes including among Vanuatu islands, continental silvereyes, and recently colonised populations.

Another clade where low support was particularly noticeable was among mainland subspecies and recent colonisations in the ANZO group where some relationships were fully unresolved. Heron Island *Z. l. chlorocephalus* and Queensland *Z. l. cornwalli* individuals were either: part of the same clade (IQ-TREE), formed a polytomy (SVDquartets, Fig. S4), or Heron Island emerged as sister to all mainland subspecies and recent coloni sations (SNAPPER) (Fig. 1B), with a divergence age of approximately 80,000 years ago (95% CI=0.53-1.13) (Fig. S5). The mainland populations and recent colonisations showed differences depending on the method used but some patterns emerged repeatedly: South and Western Australia were a single cluster, the North Island of New Zealand, Norfolk Island and French Polynesia often grouped together, as well as Chatham Island and the South Island of New Zealand. Finally, silvereyes from Tasmania and the Eastern coast of Australia (Victoria and New South Wales) were part of the same clade.

### Population structure: genomic and morphological data

We visualised genetic population structure by plotting the two PCA axes that explained most genomic variation and the results from the admixture analysis. We repeated each analysis using three datasets: i) the entire sample set including all silvereyes and white-eye outgroups, ii) a subset only including silvereye samples from the ANZO group, and iii) a subset including only samples from Southern Melanesia. Together, these analyses supported the existence of three major groups: Southern Melanesian silvereyes, ANZO silvereyes and other *Zosterops* species. The first principal component in the PCA of the entire sample set explained 38.4% of the variation, and separated Southern Melanesian silvereyes from all other silvereye populations and other *Zosterops* species, while the second (20.9%) separated all silvereyes from other *Zosterops* species (Fig. 2A). Within Southern Melanesia, the silvereyes from the New Caledonian archipelago (Grand Terre and the Loyalty Islands) were distinct from those from North and Central Vanuatu islands, with Efate and Tanna populations intermediate between the two archipelagos (Fig. 2A). These groupings, along with the distinctiveness of Tanna and Efate, were supported by admixture plots (Fig. 2B). Assignment tests showed that individuals from Tanna, Efate, Gaua, Espiritu Santo, and Pentecost were correctly assigned to their original populations (Table S3). In contrast, individuals sampled from populations in the Central and Northern Vanuatu region were incorrectly assigned to Pentecost. Additionally, we observed a significant level of structure within the New Caledonian archipelago.

**Fig. 2.**
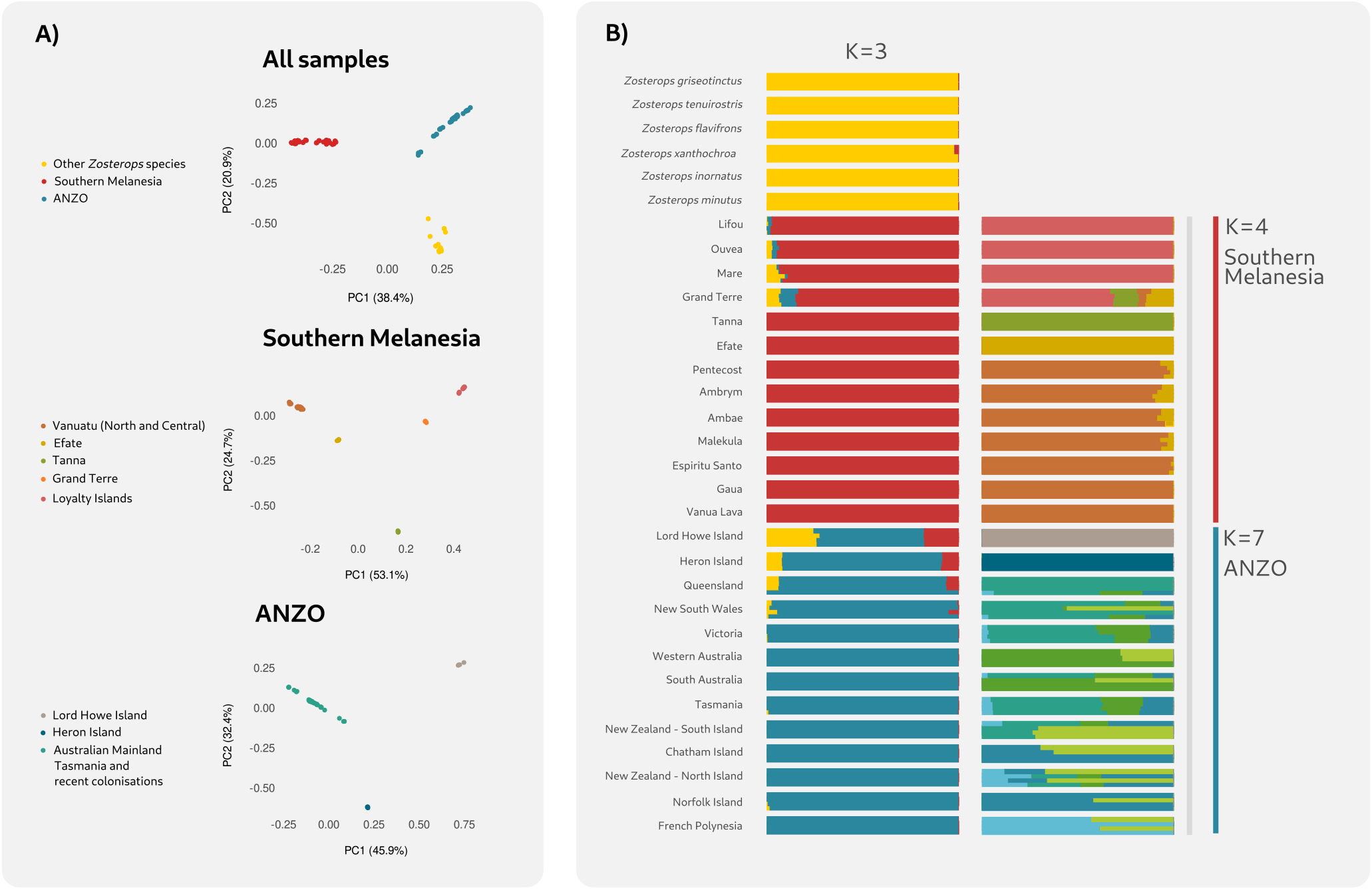
Population structure of the silvereye clade and other South Pacific white-eyes. A) Genomic PCA plots representing the two axes that explain the most variation (PC1 on the x-axis and PC2 on the y-axis) for the three datasets detailed in the methods section. B) NGSadmix bar plots for: all samples (*K* =3), Southern Melanesia (*K* =4), and ANZO (*K* =7). The represented *K* is the most likely number of ancestral populations. Each bar is the estimate of the individual’s ancestry proportion from each of the assumed ancestral populations.

Within the ANZO group, the PCA revealed three distinct clusters: Lord Howe Island, Heron Island, and a larger cluster comprising Australian mainland samples, Tasmania, and recently colonised populations (Fig. 2A). This was also supported by the admixture analysis, with the historically documented colonisations showing some finer substructure (Fig. 2B) that was also evident in assignment tests. The main patterns to note were: Western Australia and South Australia were part of the same cluster; overlapping genetic profiles among widely separated eastern Australian sites (Queensland, New South Wales, and Victoria), and the continental island of Tasmania; and the recently colonised populations showed a similar genetic makeup, except for Norfolk Island and French Polynesia where individuals were mostly classified under their own cluster.

Training a Random Forest classifier on morphological data revealed that some island populations, such as those from Tanna, Lord Howe, and Grand Terre, could be accurately distinguished from other silvereye populations based on their morphology alone (Fig. S3). Other Southern Melanesian populations sometimes clustered together, especially across Central Vanuatu (Malekula, Espiritu Santo and Pentecost), and two of the Loyalty Islands (Lifou and Ouvea). Heron Island was clustered with islands from Vanuatu despite not being close genetically. This result is not surprising as island populations frequently experience parallel evolution of body size, a phenomenon known as the ‘island rule’. In line with the genetic results, it was harder to morphologically distinguish among recently diverged populations and among populations sampled on the Australian mainland.

### Network analyses, admixture and gene flow

The Neighbor-Net agreed with other methods on the main split between other *Zosterops* species, Southern Melanesian silvereyes and the ANZO group (Fig. 3A). However, Lord Howe Island emerged as an independent lineage closer to other *Zosterops* species than to silvereyes. When we analysed the ANZO subset in detail, which included Lord Howe Island, we found that the samples from this island, Heron Island and Queensland showed mild reticulation at the base (Fig. 3C). Reticulation was more evident in samples from South and Western Australia, and Victoria and Tasmania, a pattern consistent with extensive gene flow across mainland populations, as supported by the Treemix results (Fig. 3C). In Southern Melanesia, islands in New Caledonia showed clear independent lineages divided by island, with reticulation being evident between Ouvea and Lifou, and a bit weaker between Mare and Grand Terre (Fig. 3B). Similarly, Tanna and Efate showed some reticulation among but were primarily independent lineages. Central Vanuatu islands did not show any evident clustering, but Northern Vanuatu islands (Gaua and Vanua Lava) did.

**Fig. 3.**
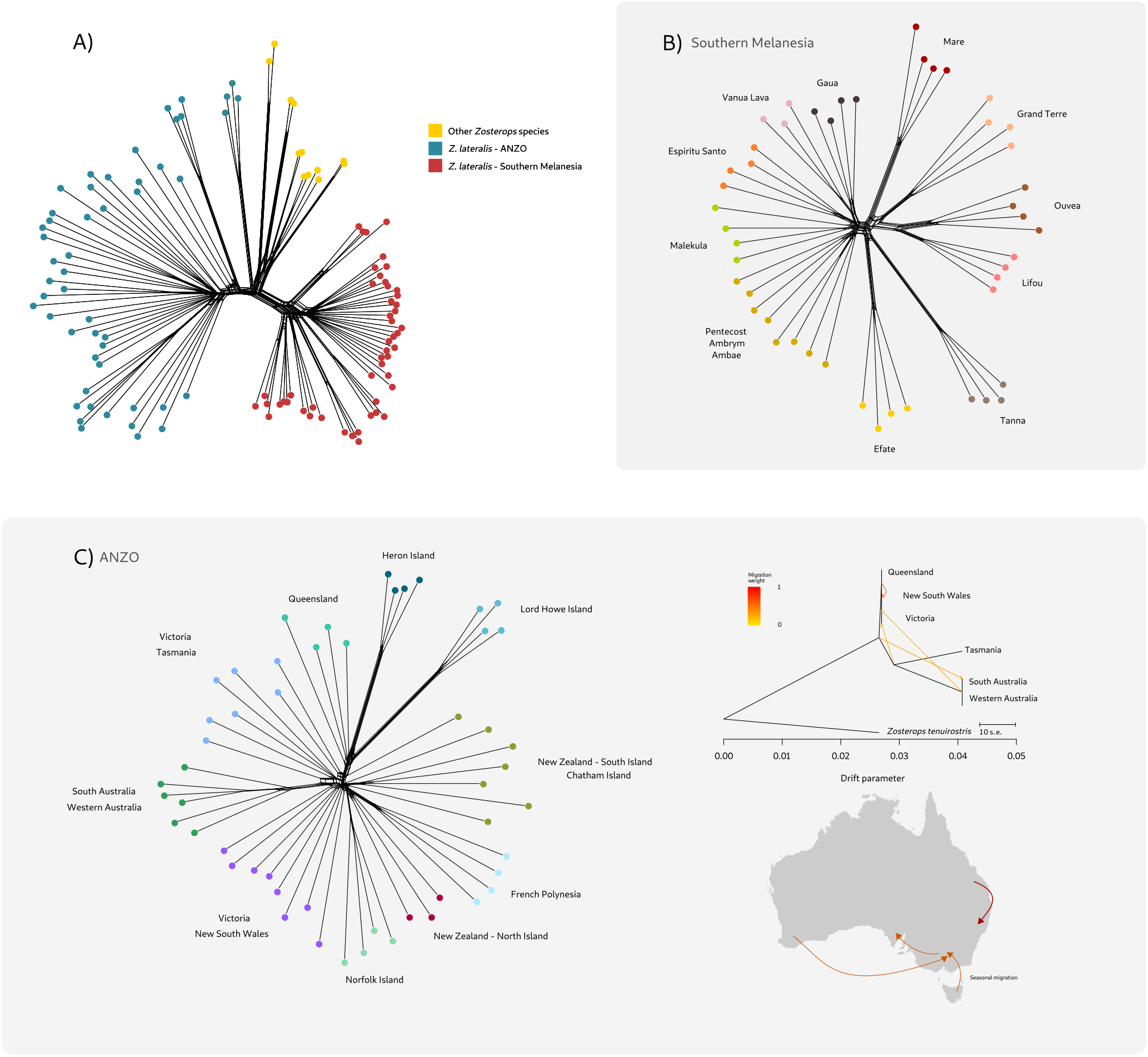
Phylogenetic network representing the evolutionary relationships and reticulate evolution. A) Silvereyes and other South Pacific *Zosterops* species, B) Southern Melanesian silvereyes and C) silvereyes from the ANZO cluster, which include continental species that show little clustering, partly due to high levels of gene flow as shown in the TreeMix maximum-likelihood tree on the right. The main directions of gene flow are shown on the map of Australia. Colours represent different populations.

#### Water barriers but not continental distances promote population differentiation

We explored whether water barriers had a stronger impact on population differentiation than continental distances, and found that F_ST_ was greater between insular populations than between continental populations (Fig. 4A; marginal effect at the mean = -0.88; 95% CI [-1.8, -0.034]; Posterior probability = 0.95). F_ST_ increased with distance on islands but not on the continent, although the latter showed wide credibility intervals (Fig. 4B).

**Fig. 4.**
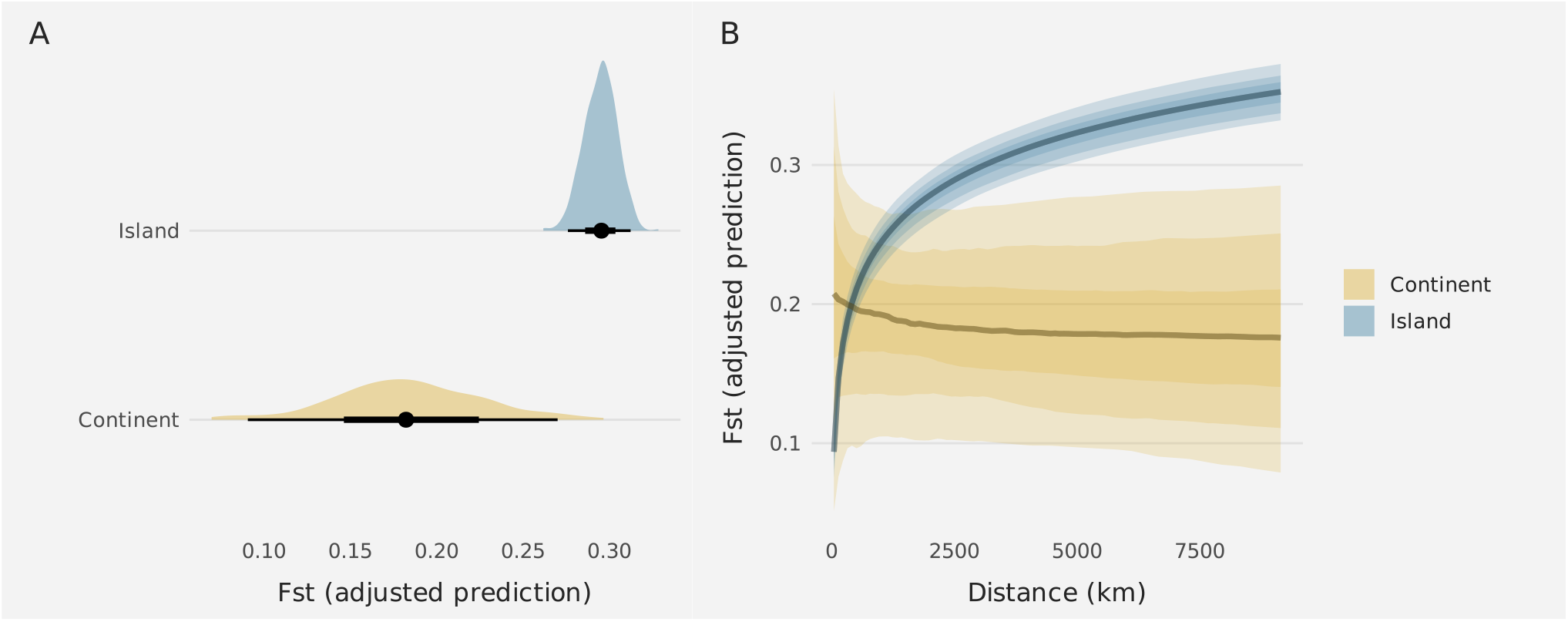
Population differentiation is stronger between islands. A) Adjusted predictions of F_ST_ on islands and the continent. B) Adjusted predictions of the effect of the distance on F_ST_ levels among continental populations (blue) and islands (yellow). The uncertainty bands are darker for lower values and lighter for higher values, representing 50% 80%, and 95% credibility intervals.

## Discussion

Silvereyes are excellent island colonisers and have overcome long-distance water barriers multiple times. Given their high dispersal ability, we would expect little population structure. This is the case in continental silvereyes, where despite long distances and ancient biogeographic barriers they show very high levels of connectivity. However, on islands we found a pronounced structure; even geographically close populations were often highly distinct, suggesting that water gaps can be significant barriers even over short distances. Overall these results indicate that islands play a central role in the diversification of a great speciator.

### The evolutionary history of South Pacific white-eyes

We first discuss the broad evolutionary relationships among the white-eye species in our dataset to provide the framework for resolving finer scale patterns within the silvereye. Our analyses agreed that the silvereye is a monophyletic clade sister to other South Pacific *Zosterops* species and that this divergence event dates approximately 1.5 Ma, during the Pleistocene, when this genus was rapidly colonising a large geographic area throughout the tropics and subtropics, from Asia to sub-Saharan Africa and Oceania (Gwee et al., 2020; Moyle et al., 2009). Our divergence time estimate is very close to both the divergence between the Afrotropical and Australasian *Zosterops*, and *Z. lateralis* from other *Zosterops* species (Martins et al., 2020; Vinciguerra et al., 2023; Wickramasinghe et al., 2017). The overlap in divergence times can be explained by the fact that white-eyes show one of the fastest diversification rates across all vertebrates (Moyle et al. 2009) thanks to their high dispersal ability, colonisation success and subsequent rapid morphological differentiation (Sendell-Price, Ruegg, & Clegg, 2020), and resilience against extinction. Their dispersal ability is especially important for understanding the mechanisms underlying speciation on islands. For example, high levels of dispersal can lead to multiple colonisation events at different points in time leading to species living in sympatry. This idea could explain why two species endemic to Lifou (*Z. minutus* and *Z. inornatus*) are not part of a monophyletic clade. These two species differ in a series of characteristics, with body size being the most striking one: *Z. inornatus* is among the largest of white-eyes (23 g) and *Z. minutus* the smallest (8 g) (Cornetti et al., 2015). According to our phylogeny, *Z. inornatus* must have colonised the island earlier, evolving a large body size as an adaptation to island life, a common pattern seen across *Zosterops* (Estandía, Sendell-Price, Robertson, et al., 2023), and then rapid evolution towards small body sizes via character displacement could have allowed a later arrival to co-exist (Black, 2010; Grant & Grant, 2006). While it is clear that the two species of Lifou white-eye are not closely related, other relationships across the non-silvereye lineage are less clear. For example, *Z. tenuirostris*, endemic to Norfolk Island, appeared both as basal to the New Caledonia species and as sister to *Z. inornatus* in different analyses. Some species of this genus are known to hybridise (Gwee et al., 2020; Manthey et al., 2020) which includes potential admixture across Norfolk Island *Zosterops* (Gill, 1970). If this were the case, gene flow across these species could explain the uncertainty at this node, but further evidence with modelling that accounts for fluctuations in population size is needed.

### Phylogenetic relationships and colonisation routes on the silvereye

The silvereye’s likely place of origin is thought to be the Australian mainland, serving as the ancestral source from which all island lineages subsequently emerged, with one of the earliest colonisation events being to Southern Melanesia. Typically, colonisation follows a one-way pattern from continents to islands, although there are exceptions, like the recolonisation of the mainland by *Anolis* lizards (Patton et al., 2021). In the south-west Pacific, colonisation is expected to mainly occur eastward from the larger landmasses, like Australia. The common ancestor could have colonised the mainland through the New Guinea-Australia land bridge that was only covered by the ocean 10,000 years ago (Friedlaender et al., 2008). However, other robust white-eye phylogenies support a more distant relationship between the silvereye and the New Guinea white-eye *Z. novaeguineae*, and a closer relationship with other island species, like *Z. flavifrons* or *Z. explorator*, endemic to Southern Melanesia and Fiji respectively (Oliveros et al., 2021). This evidence raises the possibility that the Australian mainland lineage might have originated from a colonisation event originating in Southern Melanesia or other Pacific islands. However, it is worth noting that the true sister lineage to the silvereye might be extinct, and if it once existed with a distribution overlapping New Guinea and Australia, it could lend support to the hypothesis that silvereyes colonised Australia via the New Guinea-Australia land bridge.

Southern Melanesian silvereyes showed complex phylogenetic patterns, possibly exacerbated by high levels of ILS and gene flow, which can be better represented by phylogenetic networks that allow for the representation of reticulate evolution. We found extensive reticulation across Ouvea and Lifou, aligning with their high morphological similarity. In contrast, Grand Terre and Mare showed less reticulation and were morphologically distinct despite the short distance that separates them and the other Loyalty Islands. Interestingly, Ouvea and Mare silvereyes are classified under the subspecies *Z. l. nigrescens* despite Lifou being in the middle, separating the other two islands and having its own endemic subspecies, *Z. l. melanops*. Visual distances in the PCA capture the geographic distances across Southern Vanuatu islands, as expected under isolation by distance. The most isolated of these islands, Tanna, is also home to the most genetically and morphologically distinct population. In contrast, islands across Central and Northern Vanuatu show very little differentiation, likely due to high levels of gene flow (Clegg & Phillimore, 2010; Estandía, Sendell-Price, Oatley, et al., 2023). The current subspecific taxonomic divisions in silvereyes from Vanuatu do not reflect the genetic structure that we infer here. There are two currently accepted sub-species: *Z. l. vatensis* on Tanna, Efate, Malekula, and Ambrym, and *Z. l. tropica* on Pentecost, Ambae, Espiritu Santo, Gaua and Vanua Lava, even though there is no extensive population differentiation across the archipelago with the exception of Tanna and Efate. However, subspecific taxonomy is complicated and often relies on plumage patterns (Winker, 2021) that are not represented in our analyses.

Lord Howe Island silvereyes *Z. l. tephropleurus* split from the mainland lineage approximately 1 million years ago. This subspecies seems to have been isolated from other silvereye populations as there is no reticulation with any of them. Additionally, individuals from this population have a fully distinct morphology. Interestingly, in the species network, Lord Howe Island shows very mild reticulation with other *Zosterops* species, being closer to them than to either of the two silvereye clades. Possible admixture could have occurred between *Z. l. tephropleurus* and other South Pacific white-eyes, including the robust white-eye *Z. strenuus*, an endemic white-eye species to Lord Howe Island that became extinct in the early 20th century after the introduction of rats (Day, 1981).

*Z. l. tephropleurus* is sometimes treated as a full species, however, this would make the silvereye a paraphyletic species by not containing all of its descendants.

Our divergence estimate for Heron Island is much older than we expected (80,000 years ago versus 4,000 that we set as a prior, representing the earliest time that Heron Island was vegetated). Our Bayesian analyses agreed that Heron Island was sister to the rest of the mainland subspecies, but pushed this node too far back. We think that these results are more likely due to an artefact and it can be potentially explained by the combination of the use of nuclear markers instead of mitochondrial markers (e.g. *cytochrome-b*) for which we know the rate of molecular evolution (Arcones et al., 2021), and the strict clock model we used as imposed by SNAPPER. Heron Island was the third island to be colonised from the mainland and despite the fact that many studies have not detected any gene flow (Clegg et al., 2002; Degnan, 1993; Estandía, Sendell-Price, Oatley, et al., 2023; but see Sendell-Price, Ruegg, Anderson, et al., 2020), we find patterns that indicate potential low levels of admixture including very mild reticulation between samples from Queensland and Heron Island. Depending on the method used, Heron Island samples were sister to all mainland samples and recent colonisations, or sister to the Queensland samples. This conflict could be explained by gene flow between both populations. There are reports of mainland silvereyes arriving on Heron Island (Clegg pers. obs.), but they do not necessarily breed.

Mainland birds are indeed highly dispersive: we find extensive connectivity across the eastern coast of Australia, where all populations have a similar genetic makeup, and individuals cannot be confidently assigned to their original locations. Additionally, there is evidence of strong gene flow between Queensland and New South Wales. Our results are consistent with what we know about silvereye movements in this region since the Tasmanian silvereye and population of *Z. l. westernensis* from Victoria display partial migration, whereby many individuals migrate during winter, a number of which reach southern Queensland (Chan, 1995; Mees, 1969). Tasmania is a continental island that was connected with the mainland by a land bridge until the end of the Pleistocene, approximately 12,000 years ago (Kennett et al., 2018). Despite this barrier, Tasmanian silvereyes continue to display partial migration, which is triggered by changes in light during winter (Chan, 1994). Silvereyes from South Australia share a similar climatic niche, which could potentially trigger migration, but the habitats available northwards are not suitable. Individuals from Western and South Australia did not show any differentiation despite being sampled more than 2,000 km apart and having to overcome substantial biogeographic barriers - the Nullarbor Plain and the Eyrean barrier. At the same time, samples from Victoria and South Australia, collected less than 1,000 km apart, showed some evidence of admixture but clear differentiation. The mechanisms leading to differentiated silvereye mainland populations do not seem to always coincide with geography alone and other mechanisms could be playing a role in determining genetic structure.

We know that the most recent colonisations stem from Tasmania because the arrival of silvereyes to New Zealand is historically documented and the subspecies that arrived is *Z. l. lateralis*, endemic to Tasmania (Mees, 1969). Since the arrival on the South Island of New Zealand, silvereyes colonised Chatham Islands, the North Island of New Zealand and Norfolk Island (Mees, 1969). The French Polynesian populations are a result of a human-assisted introduction consisting of a low number of silvereyes from the South Island of New Zealand in 1937 (Guild, 1938). Our results could not accurately infer the historical trail because this range expansion happened in less than 100 years, suggesting extremely short periods of time between divergence events and high ILS. Similarly, morphological characteristics were too little to be able to distinguish among recently colonised populations, which is likely due to the short amount of time to accumulate enough differences (but see Sendell-Price et al., 2020a). However, we do see in the admixture plot that the genetic makeup of late colonisations resulting from several bottlenecks are less diverse, reflecting the loss of alleles as expected in founder effects (Clegg et al., 2002; Sendell-Price et al., 2021).

### Insular population differentiation and the paradox of the great speciators

Water gaps serve as important barriers for dispersal and gene flow among silvereye populations, leading to higher genetic differentiation. This is not the case for continental populations where, despite long distances and geographical barriers, we see very little genetic structure and no effect of distance on F_ST_ levels. This pattern could be explained in different ways. For example, if island colonisation is rare but there are a few events of successful population establishment, isolated populations could arise that would eventually become separate lineages. However, connectivity patterns on the mainland suggest silvereyes do have an effective ability to disperse. Additionally, overwater dispersal is not entirely rare, as mainland silvereyes regularly reach Heron Island, an island that cannot be seen from the mainland. A more plausible explanation invokes strong selection against dispersal after island colonisation leading to population divergence (Diamond et al. 1976). Individuals on island populations may retain the ability to fly long distances but exhibit a behavioural shift whereby they are reluctant to disperse overwater, a phenomenon known as ‘behavioural flightlessness’ (Diamond, 1981). This idea fits with previous results that indicate that older and more sedentary populations have experienced changes in dispersal genes when compared with mainland and recently colonised populations (Estandía, Sendell-Price, Oatley, et al., 2023). We see a sharp increase in between-island F_ST_ with distance, suggesting that even short gaps of water can be effective barriers. This is evidenced across southern Vanuatu and the Loyalty Islands where nearby islands show fully genetically distinct populations. However, there are a few exceptions across central Vanuatu where there is evidence of reticulate evolution. In this region, silvereyes have likely been able to maintain populations connected, but there is no obvious reason why. Spatial processes, such as how islands are geographically arranged, could be a third explanation for the observed patterns. This argument does not necessarily require invoking selection against dispersal ability, and it could easily arise with intermediate dispersal (Ashby et al., 2020; Yamaguchi, 2022). Most likely a great speciator pattern would arise from a combination of spatial, abiotic and biotic factors, but the relative importance of each factor is currently unknown. To solve this evolutionary paradox, we should take different approaches, such as theoretical, by building models that help us understand the important parameters, comparative, by exploring the factors driving divergence in numerous great speciators, and empirical, by examining whether island populations show morphological, behavioural, and genetic signatures of reduced dispersal

## Conclusion

This study used genome-wide markers to explore the evolutionary history of a great speciator and whether island and continental settings have promoted population structure and differentiation. We find that islands generally led to higher population differentiation and continental populations were highly connected despite the presence of major biogeographic barriers. We resolved the main deep relationships, but many shallow nodes are unresolved despite the use of a large genomic dataset. Phylogenetic conflict likely arises due to a combination of gene flow and high levels of ILS. Future work should focus on the specific role of gene flow in diversification by tailoring demographic models that account for bottlenecks resulting from island colonisation.

## Author contributions

Conceptualization: AE, SMC, BCR; Data Curation: AE; Formal Analysis: AE (Bioinformatics), NMR (Morphology); Funding Acquisition: BCR, SMC; Investigation: AE; Methodology: AE, NMR; Project Administration: AE, SMC; Resources: AE, ATS-P, DAP, BCR, SMC; Supervision: SMC; Visualization: AE; Writing - Original Draft: AE; Writing - Review & Editing: AE, NMR, SMC, DAP, ATS-P, BCR.

## Data and code availability

Code to reproduce all analyses is available at https://github.com/andreaestandia/0.0_phylo_silvereye https://github.com/nilomr/silvereye-morphology

## Acknowledgements

The samples used in this study were collected over two decades and we thank the many people who helped in numerous ways to facilitate the work. We are grateful to the chiefs and landholders of Vanuatu and New Caledonia for granting access to field sites, and field and logistic assistance from many people including the following: N. Clark, D. Treby, J. LeBreton, F. Cugny, W. Waheoneme, O. Hebert and A. Rouquie (New Caledonia); F. Robertson and C. Sendell-Price (Heron Island); S. Geiger, O. Boissier, R. Hills, S. Totterman, O. Drew, K. Ser, W. Ser, J. Saksak (Vanuatu), I.P.F. Owens, N. Clark (Australian mainland); J. Kikkawa (Norfolk Island); I.P.F. Owens (Lord Howe Island); P. Park, A. Fletcher, P. Gray (Tasmania); P. Schweigman, D. Onely (New Zealand); M. Bell (Chatham Island); N. Davies (Moorea, French Polynesia), O. Grant, C. Sendell-Price (Tahiti, French Polynesia). For additional samples we thank A. Phillimore, R. Black, M. Massaro, and N. Clark. The work was conducted under permits from the Direction de l’Environnement Province Sud and Direction Du Développement Economique (New Caledonia and the Loyalty Islands with thanks to G. Kakue); Vanuatu Environment Unit letters of permission and permits provided by E. Bani and we further thank D. Kalfatak and T. Tiwok for their assistance (Vanuatu); Délégations régionales à la recherche et à la technologie (French Polynesia); Lord Howe Island Board; Environment Australia (Norfolk Island); Queensland Department of Environment and Resource Management; Parks and Wildlife Service (Tasmania). New Zealand Department of Conservation Te Papa Atawhai; Australian Bird and Bat Banding Scheme project and individual permits to SMC. Ethics clearances were provided by University of Queensland ethics committee (ZOO/165/94/ARC, ZOO/520/96/ARC/PHD, ZOO/520/97/ARC/PHD) and Griffith University ethics committee (ENV/01/12/AEC, ENV/07/16/AEC, ENV/06/20/AEC, ENV/24/13/AEC) to SMC. We thank F. Robertson for conducting lab work. We thank the funders of this work: a Marsden Fund grant (UOO1410) from the New Zealand Royal Society Te Apārangi awarded to BCR and SMC to support fieldwork on mainland Australia and Heron Island portions of the molecular work; National Geographic Society Committee for Research and Exploration Grant (9383-13) to SMC to support fieldwork in New Caledonia; Natural Environment Research Council (NERC) postdoctoral fellowship to SMC to support fieldwork in Vanuatu and New Caledonia; Heredity Fieldwork Grant from the Genetics Society to ATS-P and Percy Sladen Memorial fund to SMC to support fieldwork in French Polynesia the Department of Zoology, Oxford startup to SMC to support sequencing. ATS-P was supported by a NERC studentship (NE/L002612/1); AE was supported by a NERC studentship (NE/S007474/1) and a St John’s College Graduate Scholarship. We thank Tim Barraclough, Borja Milá, and Carys Jones for commenting on the manuscript.

