## Supplementary figures for "Islands promote diversification within the silvereye clade: a phylogenomic analysis of a great speciator"

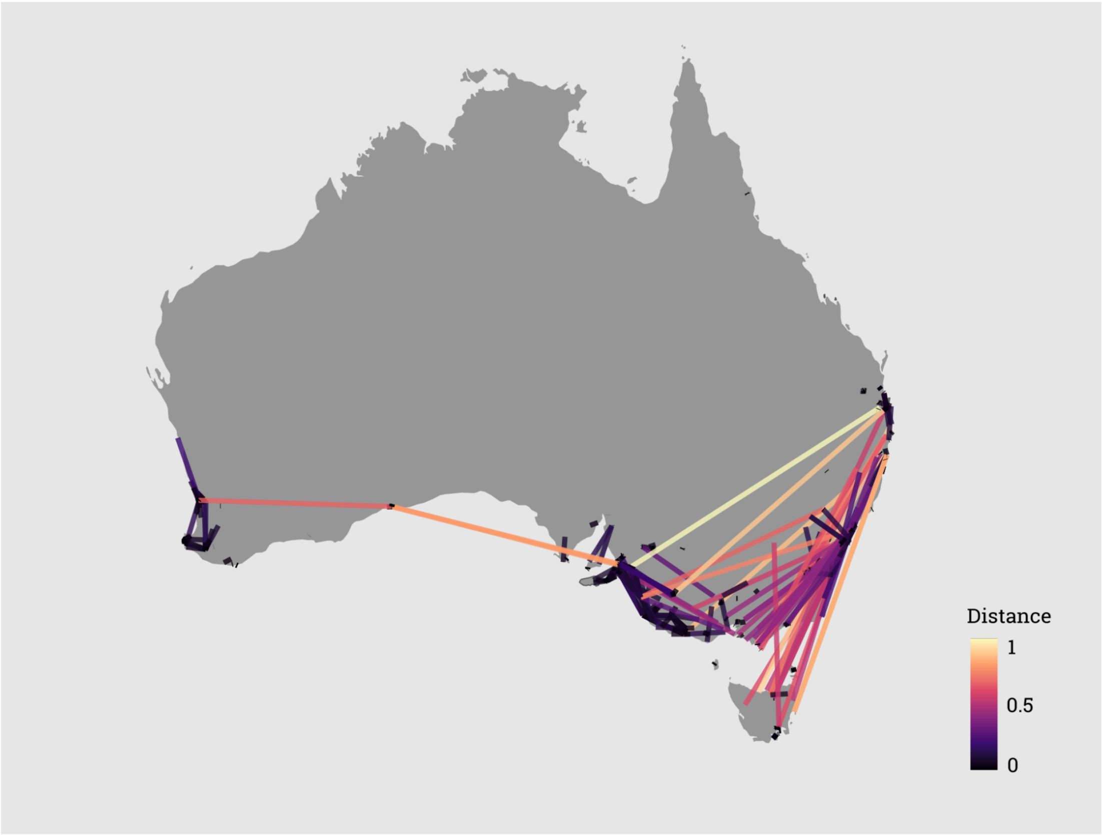


Figure S1. Silvereye movement patterns based on banding recovery records between 1956 and 2015. Lines are coloured by distance (normalised) between where the bird was first ringed and where it was caught again. Note that population density and banding effort vary greatly across space and time.

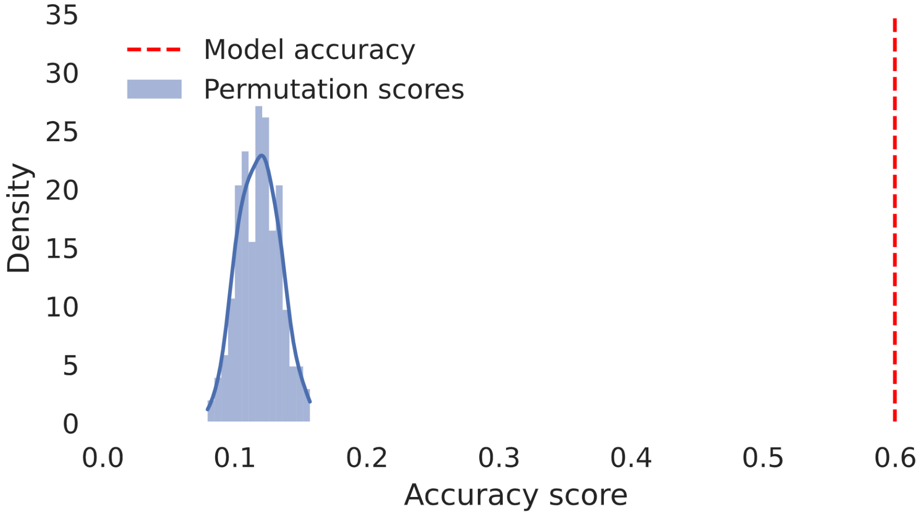


Figure S2. Permutation score test plot representing a distribution of how accurate random classification would be using morphology. The red dashed line represents our model's accuracy in predicting populations based on morphology, indicating an average classification of 60%.

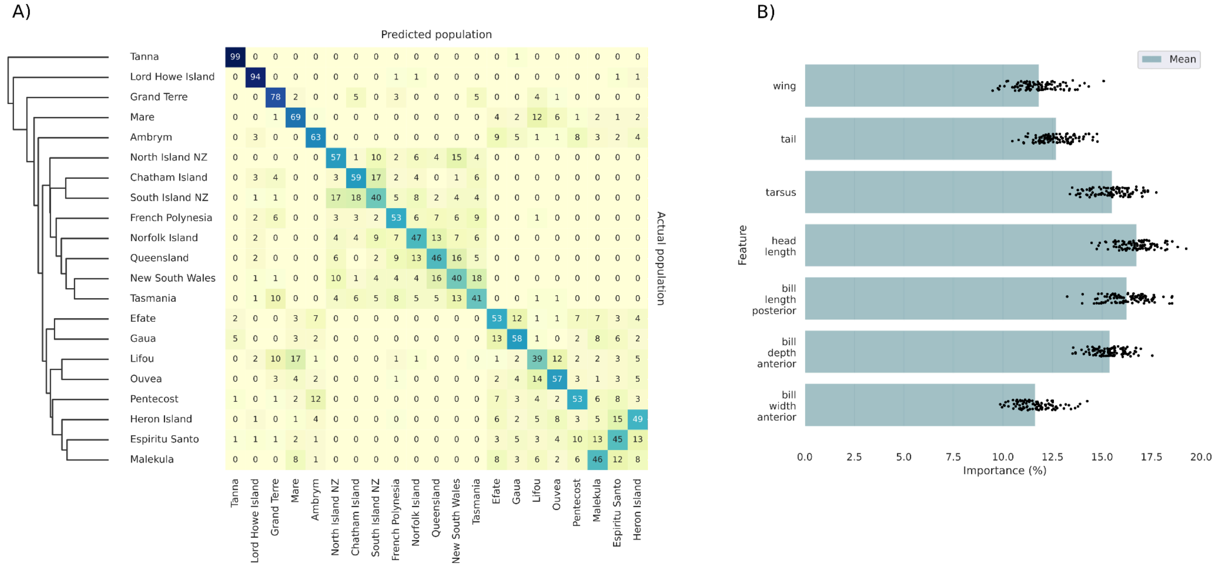


Figure S3. A) Morphological distinctiveness of populations determined by hierarchical clustering of the confusion matrix. The numbers in cells are percentages and are shaded from yellow (low percentage) to dark blue (high percentage). The diagonal shows the percentage of iterations predicted population matched the sample population and the off-diagonal values cases of mis-assignment. Tanna is highly distinctive with 99% of the island sample assigned to itself. In contrast, the value for New South Wales was 40% of the time, and it is sometimes incorrectly predicted to be from Tasmania (18%) and Queensland (16%); B) All traits are similarly important in determining the morphology.

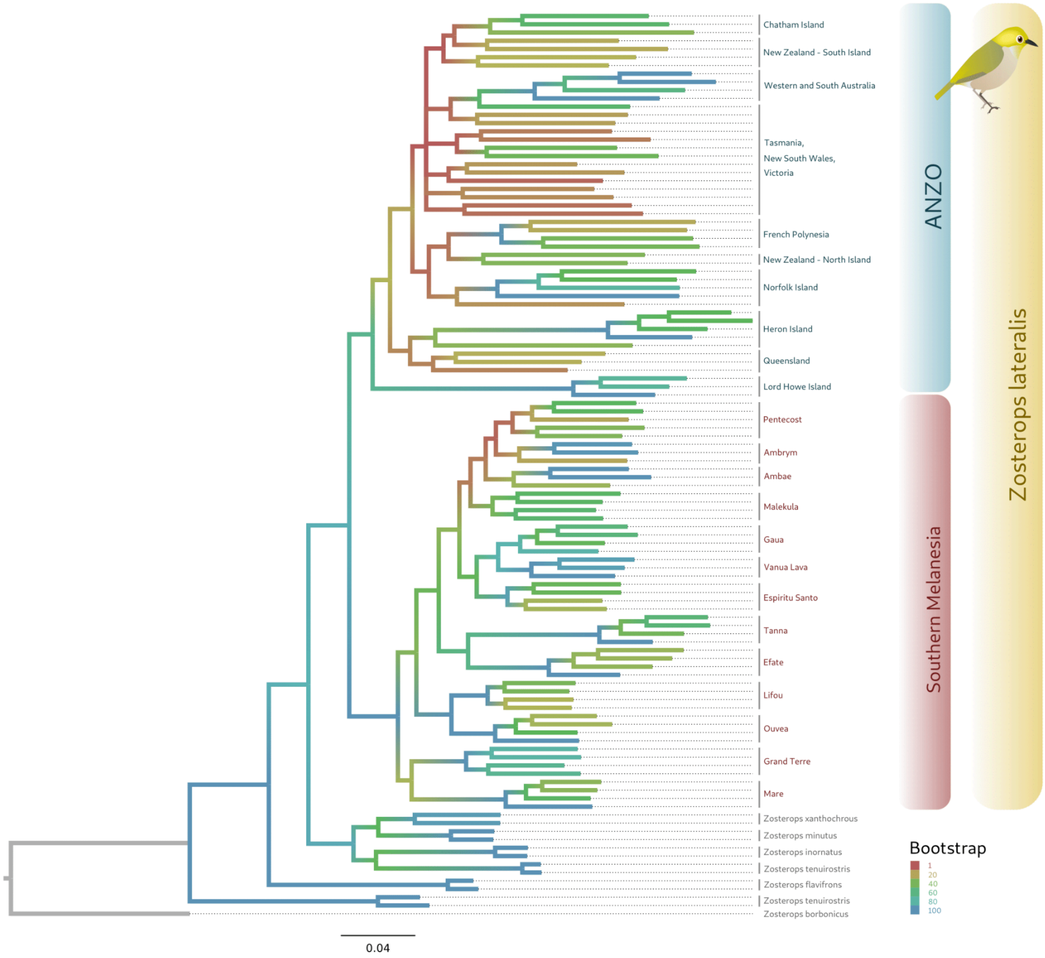


Figure S4. The phylogenetic tree resulting from the IQTREE analysis, where each branch is colour-coded to indicate the support for its parent node. This reveals significant uncertainty across the ANZO cluster (excluding Lord Howe Island) and North and Central Vanuatu.

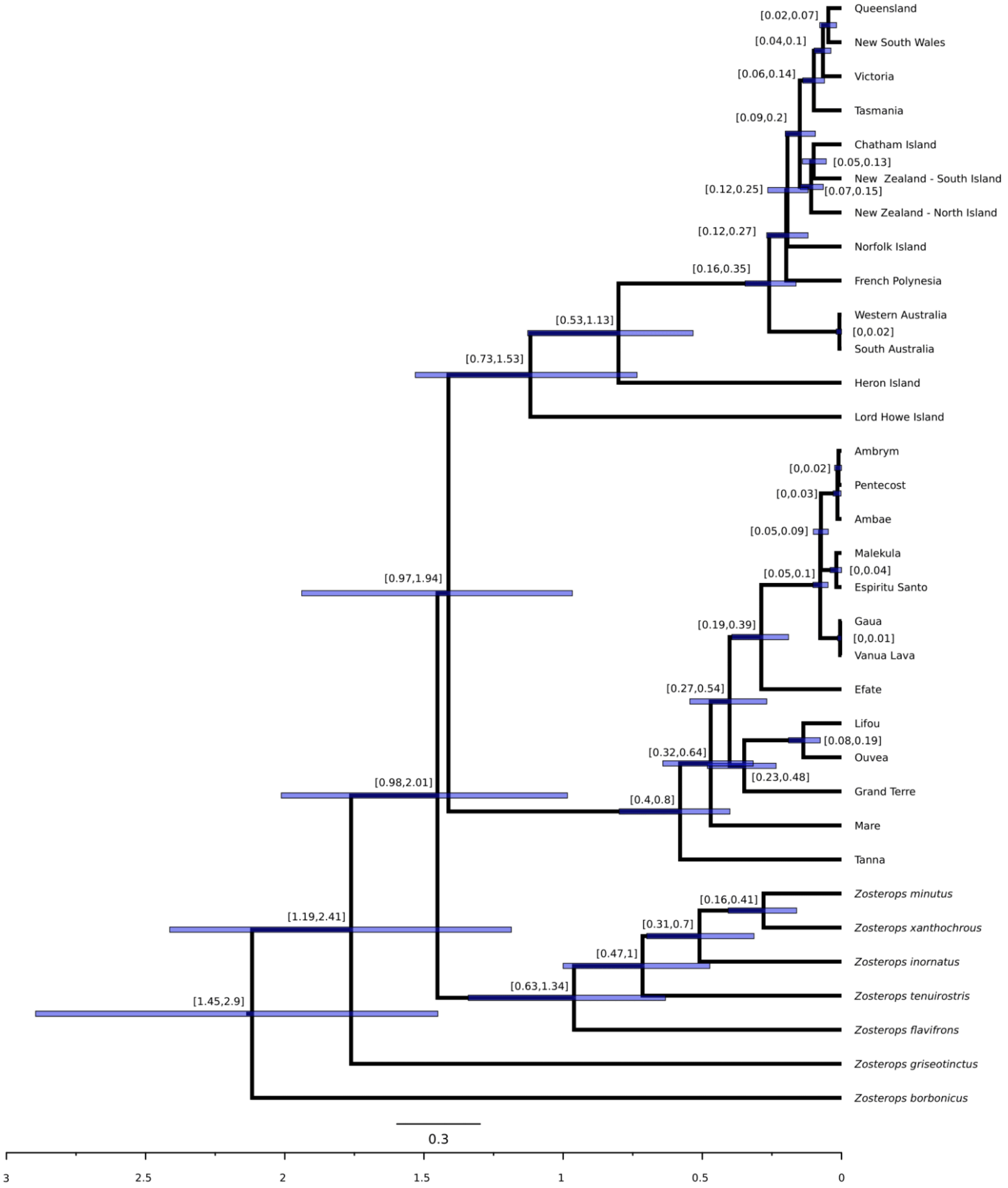


Figure S5. Maximum clade credibility tree representing the consensus topology and the 95% credibility intervals for the age estimate.

### Supplementary table captions

Table S1. Curated dataset of the samples sequenced and associated metadata.

Table S2. Curated dataset of the samples used for the morphological analysis.

Table S3. WGSassign results
